# Automatic late blight lesion recognition and severity quantification based on field imagery of diverse potato genotypes by deep learning

**DOI:** 10.1101/2020.08.27.263186

**Authors:** Junfeng Gao, Jesper Cairo Westergaard, Ea Høegh Riis Sundmark, Merethe Bagge, Erland Liljeroth, Erik Alexandersson

## Abstract

The plant pathogen *Phytophthora infestans* causes the severe disease late blight in potato, which results in a huge loss for potato production. Automatic and accurate disease lesion segmentation enables fast evaluation of disease severity and assessment of disease progress for precision crop breeding. Deep learning has gained tremendous success in computer vision tasks for image classification, object detection and semantic segmentation. To test whether we could extract late blight lesions from unstructured field environments based on high-resolution visual field images and deep learning algorithms, we collected ~500 field RGB images in a set of diverse potato genotypes with different disease severity (0-70%), resulting in 2100 cropped images. 1600 of these cropped images were used as the dataset for training deep neural networks. Finally, the developed model was tested on the 250 cropped images. The results show that the intersection over union (IoU) values of background (leaf and soil) and disease lesion classes in the test dataset are 0.996 and 0.386, respectively. Furthermore, we established a linear relationship (R^2^ = 0.655) between manual visual scores of late blight and the number of lesions at the canopy level. We also learned that imbalance weights of lesion and background classes improved segmentation performance, and that fused masks based on the majority voting of the multiple masks enhanced the correlation with the visual scores. This study demonstrates the feasibility of using deep learning algorithms for disease lesion segmentation and severity evaluation based on proximal imagery for crop resistance breeding in field environments.

## 1. Introduction

Crop disease poses a threat to global food security [1]. Automated field phenotyping can become a powerful tool for future resistance breeding as well as for precision agriculture [2][3],and can thus be a successful way to mitigate crop disease. Potato is today the third most important food crop and is an important part of many diets, especially in temperate climates. The oomycete *Phytophthora infestans* (Mont.) de Bary which causes potato late blight (PLB) and potato tuber blight (PTB) can be very destructive in potato cultivation if it is not managed (Wiik, Rosenqvist, & Liljeroth, 2018). In practice, the prevention of PLB is in the field is highly relying on regular blanket spraying of fungicide during the growth season. As an example, in Sweden the potato production consumes around 20% of all fungicides used in agriculture, largely to combat *P. infestans*, in spite of occupying less than 1% of the area under cultivation [4]. In addition, PLB prevention requires frequent use of fungicides with sometimes more than 10 applications per growth season in Northern Europe to avoid significant yield loss. This management is effective in general and widely accepted by farmers, but also results in usage of large amounts of fungicide as well as fossil fuels, which hampers the sustainable development of agriculture.

The loss caused by PLB can be reduced by breeding PLB resistant cultivars. To breed for high PLB resistance, plant breeders establish experimental plots to quantify the PLB severity of different potato genotypes and progeny lines. This is currently manually done by estimating visual scores based on the number and area of lesions on plants [4]. This process is time consuming, can be subjective and also requires experienced raters for visual scoring. Therefore, it is highly needed to develop an automated disease evaluation system to facilitate and speed up the breeding processes. However, one of the main challenges in automating the system is to accurately segment the lesions under field conditions. There are some studies which have employed PLB lesion segmentation. For example, Abdu et al [5] developed a pattern recognition approach to recognize early blight, caused by *Alternaria solani*, and PLB visual disease symptoms using soft computing and machine learning algorithms. Barbedo [6] and Camargo et al [7] also carried out similar studies on plant disease detection and segmentation. However, the image datasets used in these studies above are collected at foliage level under relatively clear and uniform backgrounds in controlled settings and the pipelines are not readily implemented into large scale field environments. Moreover, the majority of previous works have only investigated the symptom segmentation problems based on a single potato cultivar. These models might fail to segment other lesions due to differences in lesion morphology and leaf color between different potato cultivars.

Attempts have also been made for early detection of PLB. Various sensors from imaging to nonimaging sensors have been employed in these applications. For example, Fernández et al [8] investigated the classification accuracy changes of infested and healthy potato leaves over different days post-inoculation (DPI) with a spectroradiometer and a multispectral camera under structured environments. Appeltans et al [9] discussed the imaging parameter settings for hyperspectral and thermal proximal disease sensing in potato and leek fields with a ground-based vehicle. Other than ground-based platforms, an unmanned aerial vehicle (UAV) platform equipped with imaging sensors also has showed its feasibility on field PLB monitoring with the detection of spectral changes in crop traits [10] [11].

Deep learning has proved to be an effective approach for traditional computer vision problems such as image classification, object detection and segmentation, as it is capable of extracting features hierarchically [12]. In addition, the applications with deep learning in the agriculture domain also show unprecedented advancements. Specifically, in precision farming it has been deployed for weed detection [13], agricultural pest detection [14] and selective fruit harvesting [15], as well as leaf counting [16]. Furthermore, deep convolutional neural networks, one of the most used deep learning algorithms, combined with computer vision techniques have been exploited for crop disease classification and detection. Polder et al [17] adapted a fully convolutional neural network (FCN) for potato virus Y (PVY) detection based on hyperspectral imagery. It proved that the deep learningbased approach outperformed the, conventional disease assessment and indicated the suitability of this method for real-world disease detection. Stewart et al [18] [19] [20] developed the deep neural networks for northern leaf blight (NLB) in maize from field RGB images collected from an unmanned aerial vehicle (UAV) platform. By contrast, the quantification and detection of PLB lesions have still been confined at a laboratory scale. To the best of our knowledge, the use of deep convolutional neural networks has not previously been explored for PLB lesion segmentation in diverse potato genotypes based on RGB imagery from the field.

Visual scoring in the field provides an important metric to quantify disease severity, but is prone to be biased and error can be subjected to raters. Thanks to automation, effectivity and objectiveness, sensor-based measurement, especially imaging sensors, provides potential advantages compared with visual scoring. Image-based analysis including images from RGB, multispectral and hyperspectral sensors has measured disease severity under controlled conditions [21] or based on PLACL (Percentage of Leaf Area covered by Lesions) at single leaf level [22], but is yet to demonstrate its full potential for accurate estimation in field environments [23].

The specific objectives of this study are (1) to evaluate the performances of deep convolutional neural networks for PLB lesion segmentation; (2) to determinate the optimal class weights for the classes PLB disease lesion and background (i.e. leaf and soil); (3) to fuse prediction masks at multiple scales for more accurate lesion prediction; (4) to determine the correlation between visual scoring and the number of lesions at the canopy level. The early pre-symptom PLB detection is out of the scope of this study as only RGB images were analyzed.

## 2. Material and Methods

### 2.1 Image data collection

The images were acquired with a hand-held RGB camera (Sony RX 100 iii) in nadir (+/− 5 degrees) at approximately 40 cm over each canopy. The ISO, aperture and FOV of the camera were set to 125, f/5.6 and 8.8mm, respectively. Images were acquired in full cloudy, semi-cloudy or sunny light settings. No flash was used. Especially for the semi-cloudy and sunny acquisitions, consideration was taken to ensure no additional shade was being cast onto the canopy from person or camera. No post-processing regarding color correction was performed.

The field location was outside Give, Denmark (N 55.859188, E 9.331065). Figure 1 shows a single image of the field from 100 meters above ground, at 15 Days After Infection (DAI). It is clearly seen how a large part of the trial field is decimated by PLB. The trial was set up with guard rows of Oleva (cv.) around the trial area. Each plot within the trial consisted of 4 plants in a 2×2 formation with infector rows on each side and a 50 cm gap between plots. The total trial consisted of 775 genotypes: 48 genotypes with three replicates, 59 genotypes with two replicates and 513 genotypes represented by single plots. Infector rows consisted of alternating Bintje (cv.) and Oleva (cv.).

**Figure 1.**
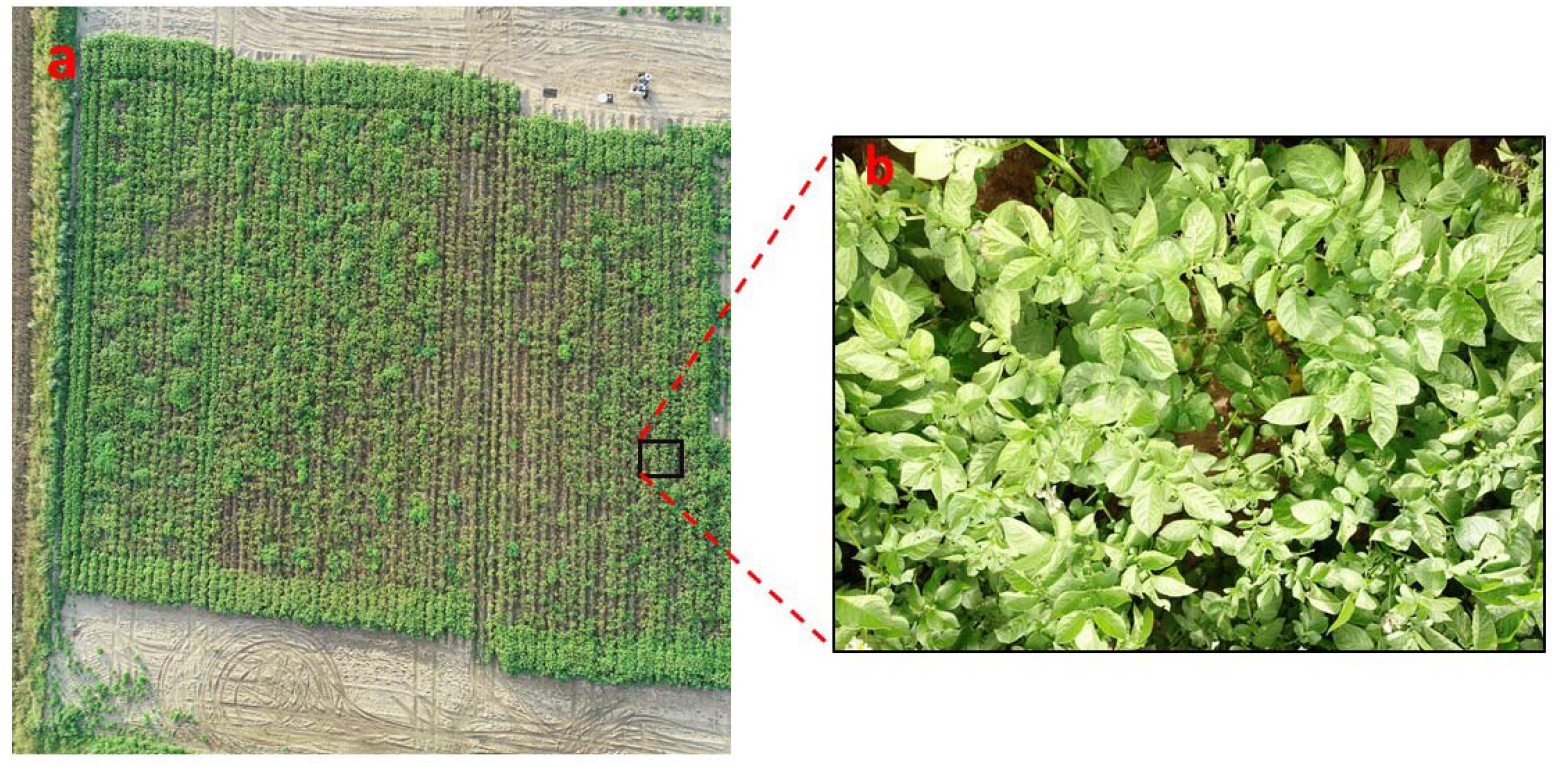
A bird’s-eye view of the experimental trial taken from a drone platform at 100 m above ground (a); an image example at 15 DAI (b).

### 2.2 Manual visual scoring of PLB

Manual visual scoring was done 2 times a week, starting 5 days after inoculation of infector rows and continuing until the standard cultivar Robijn had reached 50% infection. Infection of PLB at each time point was scored as percentage of leaf area infected according to the Euroblight protocol [24] with a single lesion per plot scored as 0.1%, 2-5 lesions scored as 0.5%, 5-10 lesions scored as 1%. The manual detection of lesions was done by walking between the two rows of each plot at a slow pace and looking at the plants at an angle. If a lesion was spotted, it was further investigated for PLB presence and the plot were investigated for presence of more lesions. Percentage of infection was used to create an area under disease progression curve (AUDPC) as follows:

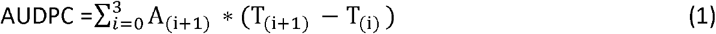

where T_0_ is innoculation date and T_i_ is evaluation dates (i=1,2,3,4). A_i+1_ is the percentage of infection at T_i+1_.

### 2.3 Image preprocessing

There are 70 original images (5472×3648) which were all labelled manually with the annotation toolbox LabelMe [25]. The distribution of genotypes among these images is shown in Figure 2. The majority of images are from the Bintje and Oleva potato cultivars. These two genotypes are susceptible to *P. infestans*, providing more disease lesions for training. The original image is unable to be directly fed to neural networks as the spatial resolution of original images is too large and requires intensive computation memories. It is also not advisable to shrink the whole image to a small size, making processing possible but heavily degrading the quality of small features in tiny lesion annotations. We first cut each original image into 6×5 (horizontal x vertical) sub-images (912×730) and then resized to 512×512 in order to feed to the neural networks for training. Each original image contributes 30 sub-images leading to 2100 sub-images in total. 1600 sub-images were randomly selected for the training set and the remaining 500 sub-images were randomly separated equally as the validation set (250 sub-images) or the test set (250 sub-images). The same procedures were applied to their corresponding ground truth images to construct image pairs for training neural networks.

**Figure 2.**
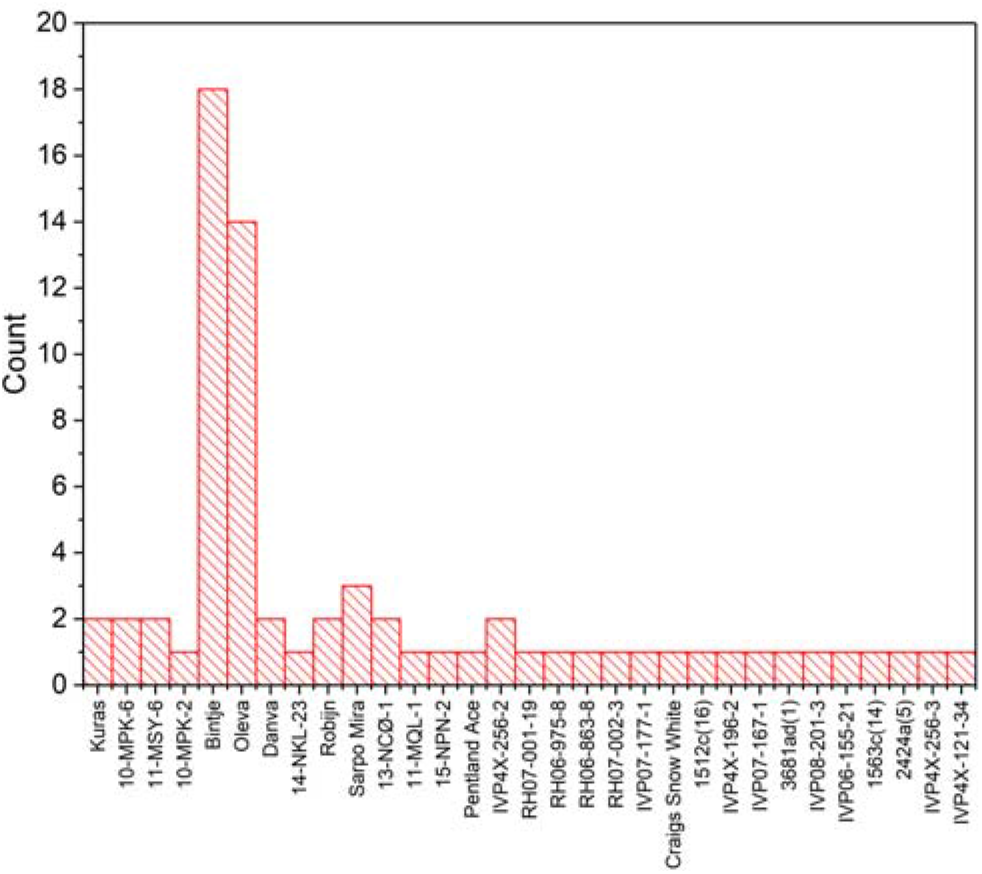
Histogram of raw image genotypes, the red bar represents the number of images belonging to that genotype.

### 2.4 Deep learning

We adopted an encoder-decoder neural network architecture based on SegNet [26] for lesion segmentation. The proposed network operates on input images of 512×512 pixels and outputs segmentation masks in the same size as the input image. The architecture has an hourglass shape consisting of a bunch of convolutional and up-convolutional layers. A diagrammatic overview of the network is shown in Figure. 3. The encoder comprises a series of convolutional operations, activation and pooling operations. Semantic features from low-level to high-level could be extracted at the end of the encoder process. Because of max-pooling layers, the spatial resolution output shrunk 2 times compared to the previous convolution block in the encoder phase. The index of the maximum feature value in each pooling window was recorded for each encoder feature map (Figure 4). In contrast, the decoder part upsamples its input feature maps using the recorded max-pooling indices learned from the corresponding layers in encoder part and finally generates the prediction results with same spatial size as original input images. The upsampling layers, such as deconvolution layers, in the decoder are also capable of learning a mapping from feature map to semantic segmentation results. Each convolutional layer is followed by several operational layers including a non-linear ReLU activation function (max(0, x)) and batch normalization. The total parameters are 29,442,122 with 29,434,694 trainable parameters and 7,428 non-trainable parameters.

**Figure 3.**
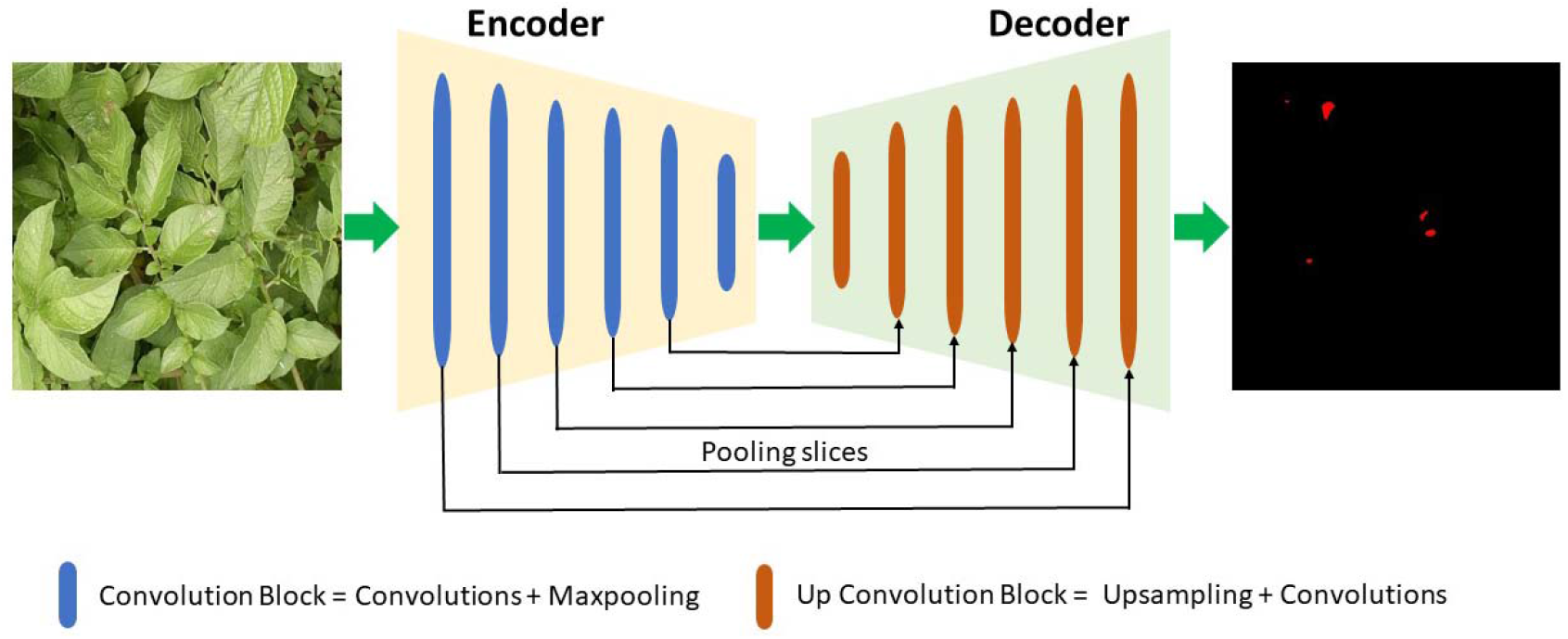
The neural network is based on an encoder and decoder architecture, followed by a final pixel-wise classification layer.

**Figure 4.**
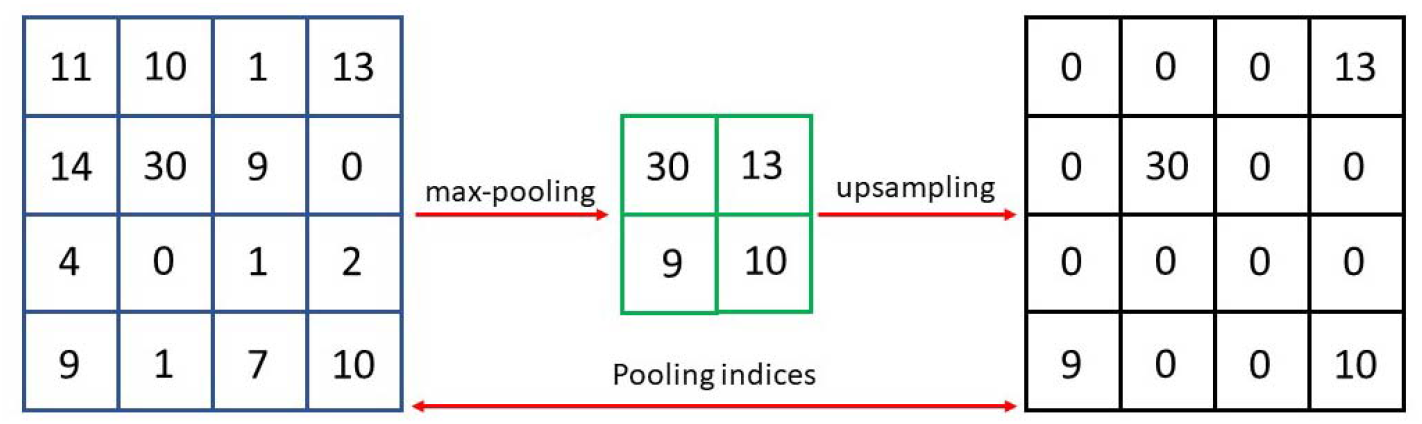
Example of max-pooling and upsampling with index in SegNet

### 2.5 Loss function

The loss function is key for training a robust and high-performance network. The most commonly used loss function for semantic segmentation is pixel-wise cross-entropy loss. In our study, the frequency of appearance for lesion and background class is highly imbalanced. The number of disease lesion pixels is far less than other pixels such as healthy plant organs and soil backgrounds in field images at early infection stages. Only using standard loss function without adaption would make a deep neural network model tend to only correctly classify dominant class pixels (backgrounds), ignoring the importance of lesion pixels. This is also called accuracy paradox, which a model provides a very high overall accuracy but performs poorly over classes. One common way to mitigate this effect is the use of a class-balancing approach by assigning different weights over classes based on their median frequency [27], In this study, we regard the weights for each class as a hyperparameter to tune. Other than weights calculated from Equation (2), we also compared the prediction performance with different weight ratios from 1 to 9 to select the optimal weight. The weighted loss function used in the network is shown in Equation (3).

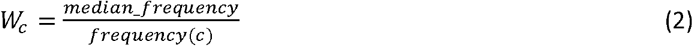

where frequency(c) represents the frequency of occurrences of pixels of class c divided by the total number of pixels in any images containing that class, and median_frequency is the median of these frequencies overall all classes.

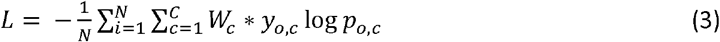

where *N* is the number of observations, *C* is the number of classes (background and lesion), *W_c_* is the weight for class *c, y* is a binary indicator (0 or 1) if a class label is correctly classified for observation *o, p* is predicted probability of observation *o* being of class *c*.

### 2.6 Training

The network was trained end to end from scratch with Adam optimizer [28] using a stable learning rate 0.0001 to minimize the loss values. The batch size was set to 18 with 500 epochs in total. We employed 3 Nvidia Tesla V100-SXM2 GPUs with around 32G memory each for training the network. Each epoch took around 171s to finish, accounting for 23.75 hours of training time in total. Data augmentation was used to reduce the risk of overfitting in the training phase. Specifically, in each batch, cropping, horizontal or vertical flipping, and a zoom range from 0.8 to 1.2 were randomly applied in the images and their corresponding ground truth masks. All network training and validation were done using the Tensorflow deep learning framework. The model was saved only with the decline of loss values in each epoch. The accuracy and loss values in the validation dataset were recorded as well in every epoch.

### 2.7 Model evaluation

The model was evaluated with three standard metrics for semantic segmentation. The three metrics are overall average accuracy, class average accuracy and mean intersection over union (mIoU), respectively. The calculations are listed below. We also used confusion matrix to check how many pixels of each class are correctly classified. Overall average accuracy (Calculation (4)) measures the performance overall all pixels. The high value means that the majority pixels are correctly classified but does not indicate good lesion segmentation as majority of pixels in our image are background. Hence, this value is sometimes quite biased for evaluating model performances. Class average accuracy (Calculation (5)) averages the performance of each class. A high value represents good performance across all classes. IoU (intersection over union), also known as Jaccard index, is a commonly used and effective metric in semantic segmentation. It measures the area of overlap between the predicted segmentation and the ground truth divided by the union area of the predicted segmentation and the ground truth in labelled images. mIoU is calculated by averaging the IoU of each class (Calculation (6)).

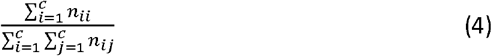

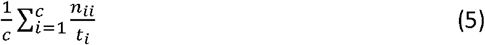

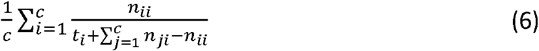

where *t_i_* is the total number of pixels of class *i* in ground truth image, *n_ji_* is the number of pixels of class *j* predicted as belonging to class *i* and *c* is the total number of classes.

### 2.8 Post-processing

Fully connected conditional random fields (FCCRFs) [29] were used first for post-processing the predicted masks. It combines single pixel prediction and shared structure through unary and pairwise terms to improve smoothness and to maximize agreement between similar neighboring pixels. The FCCRFs establish pairwise potential by using a Gaussian function on all pixel pairs in an image. The main benefits of using FCCRFs are determining the optimal decision boundary at conflict regions of pixels, while not having notable negative effects on successfully segmented pixels. In postprocessing of images, the prior knowledge of *P. infestans* disease lesion area is relatively small in an early infection stage. We found that some false positive areas likely represent shadowed area between leaves (Figure 9). These false lesion areas are far larger than normal lesions appeared at that early infection stage, so a simple threshold algorithm was operated to filter out part of false positives in the test images. We set a reasonable lesion area range to be [50, 10000] pixels to exclude the extreme false positives. Furthermore, the canopy heights and structures of potato plants vary in trial fields. That means even the same lesion spots represent differently in 2D images, which can lead to failed predictions. For some failed lesions the network can successfully predict the lesions at a different scale. A majority voting approach for lesion counting was proposed based on multiple prediction masks from various scales. The generic pseudo-code is listed in Table 2. Each image for prediction was cropped into sub-images at 7 multiple scales from 3×2 to 9×8. The sub-images with the same scale were predicted separately by the model, resulting in 7 predictions for each image. The final prediction mask of an image is obtained based on the majority voting of its 7 prediction masks.

**Table 1.**
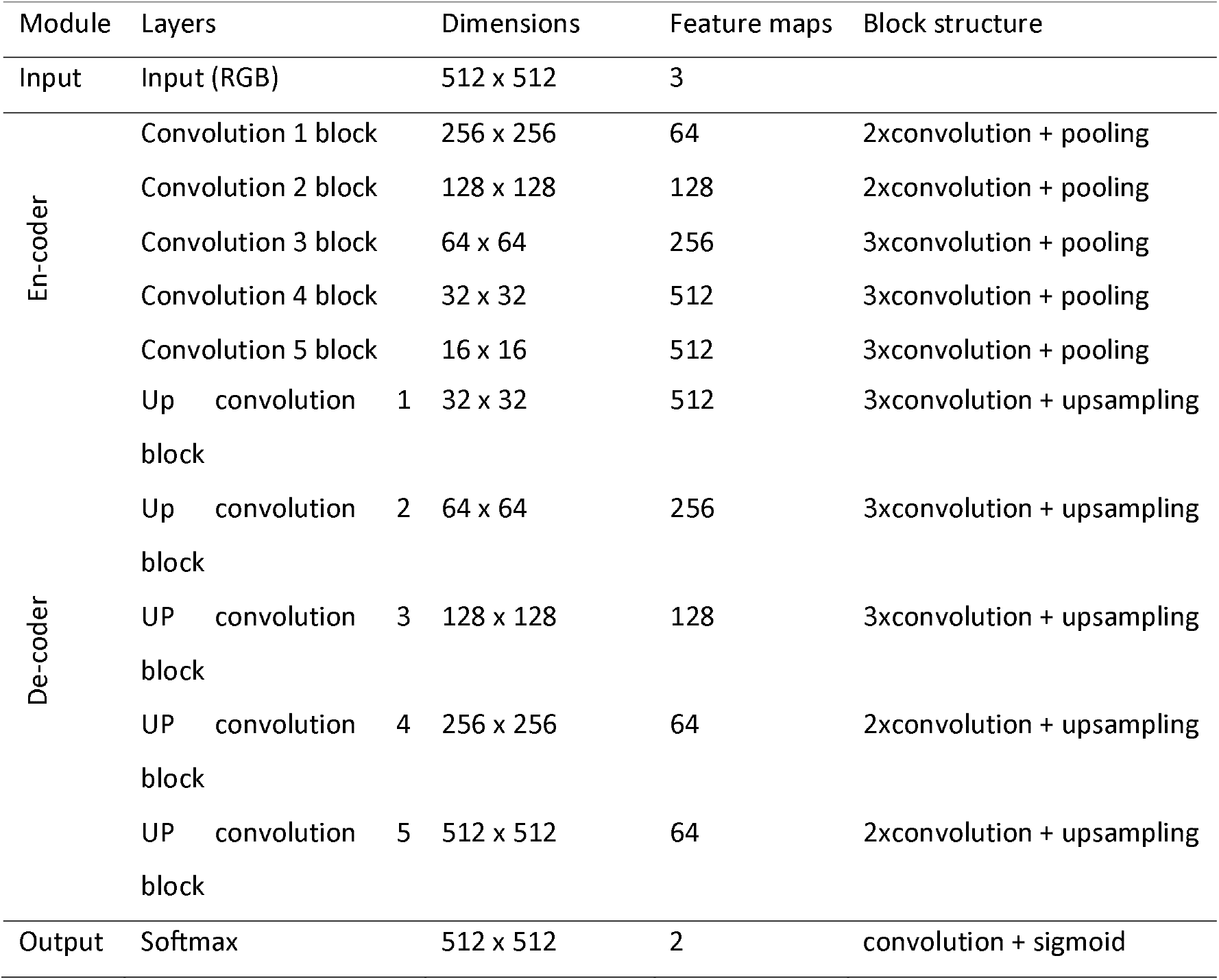
Details of the proposed deep neural network architecture. All the filter size is 3 x 3 and padding mode is set as ‘same’ with zero-filled. The final decoder output is fed to a softmax classifier to produce class probabilities for each pixel.

**Table 2.** Pseudo-codes of fused mask generation based on a majority voting approach

| **Algorithm:** Generic pseudo-code of lesion counting |  |
| --- | --- |
| 1: | **Input:** test dataset |
| 2: | **For** image in test dataset **do**: |
| 3: | **for** i=3 to 9 **do**: |
| 4: | cut image horizontally i and vertically i-1, i x (i-1) sub-images generated in total; |

## 3. Results

### 3.1 Network training

Overall, the training loss and validation loss decreased with the increment of training time (Figure 5). The validation loss fluctuated much in the early training stage (< 150 epochs) and then slowly converged at 0.0626 at the end of the training. By contrast, the training loss smoothly dropped until the end of training and finally converged at 0.0398, slightly lower than the final validation loss (0.0626). It also can be observed that both the training loss and validation loss were substantially stable after 450 epochs, indicating that the model stopped improving on a hold-out validation dataset. The model weights were saved at 450 epochs to prevent the risk of overfitting. At this epoch point, the overall accuracy values in the training and validation datasets were 0.9962 and 0.9945, respectively. The same procedures were followed when training other models with different hyperparameters (the weight ratio of lesion and background) for performance comparison in the test dataset.

**Figure 5.**
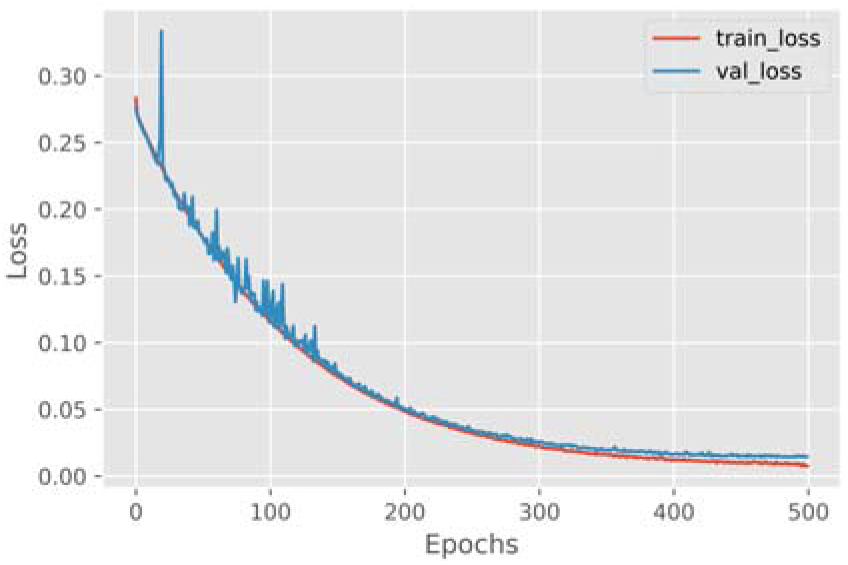
Loss curves of the network (1:7 weight ratio) in the training and validation datasets

### 3.2 The weight ratio of two classes (lesion and background)

The weight ratio of lesion and background classes is one of the important hyperparameters needed to be fine-tuned. In this study, we investigated 13 group weight ratios in order to determine the optimal weight ratio for lesion semantic segmentation. One of the weight ratios (1:2.5) was obtained based on the median frequency of two classes and the remaining weight ratios were ranging from 1 to 12. The metrics in the validation dataset are shown in Table 3. It shows that the imbalance weights can effectively improve the segmentation performance as all mIoU values exceed 0.65 compared to the 0.551 obtained when the weight ratio was set to 1:1. Interestingly, mIoU value does not continue to increase with a larger weight ratio (>7). The maximum mIoU value was achieved with 1:7 weight ratio. We selected the model with this weight ratio as the optimal model for lesion segmentation.

**Table 3.**
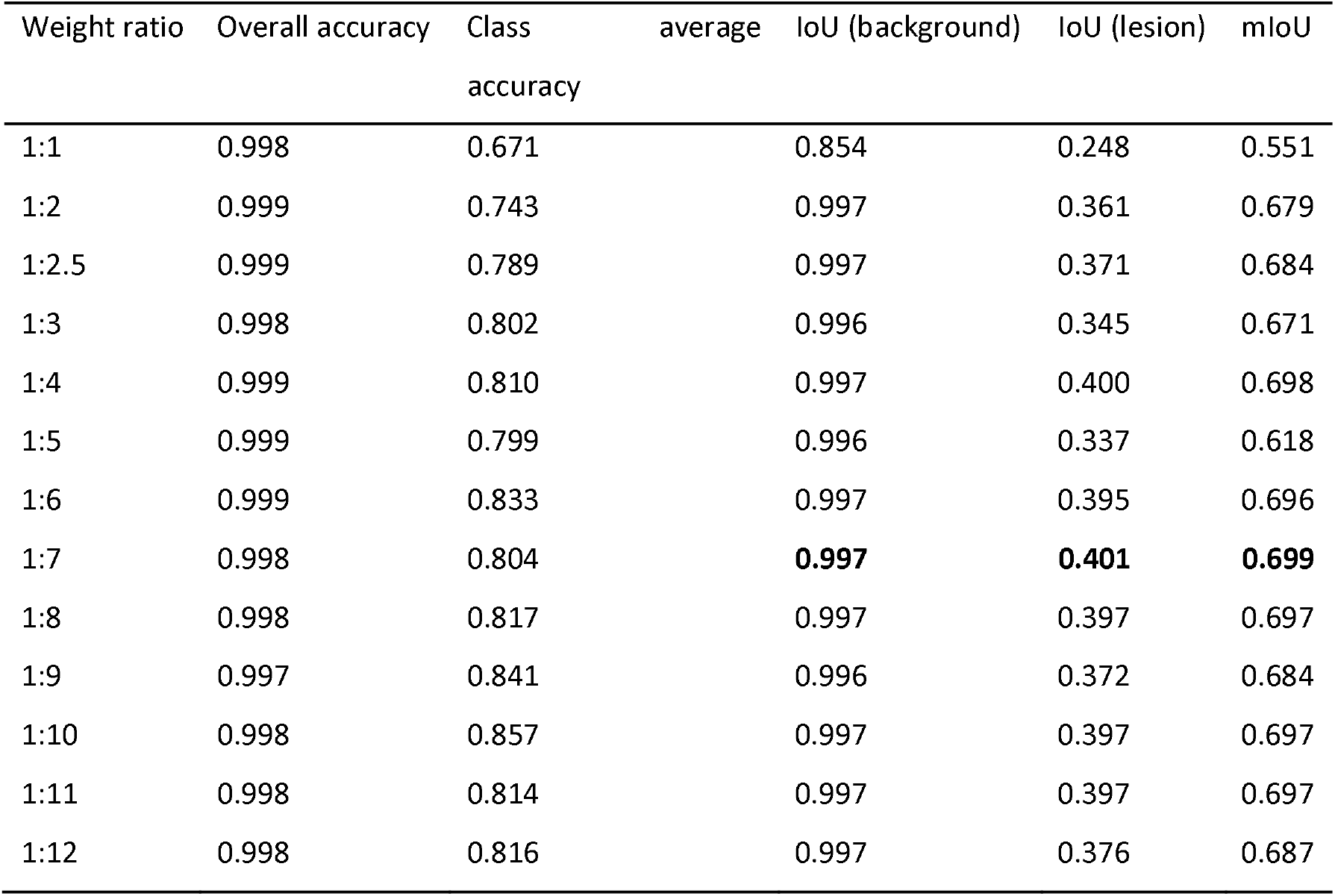
Metrics of the models with different class weight ratios in the validation dataset

### 3.3 Test image prediction

Similar to the image process for training, the original test images (5472×3648) were first cropped and then resized to sub-images (512×512) for prediction. We used the model with 1:7 weight ratio as the optimal model to test the images. Figure 6 shows confusion matrix in the validation dataset (a) and in the test dataset (b). The IoU values of background and lesion classes in the test dataset are 0.996 and 0.386, respectively. The metrics in the test dataset are lower than in the validation dataset. Most background pixels (99.8%) are correctly classified from the confusion matrix. As the majority of pixels are leaf, belonging to background class, they can be easily classified based on the color differences with lesion class. Around 40% of lesion pixels were classified as being background class in the test dataset. The prediction examples are illustrated in Figure 7. Generally, most lesions, marked as the red areas in the images, can be correctly segmented. Some tiny lesions were failed to be manually labelled on the ground truth images, but they were successfully segmented by the model (shown in #2 and #4 columns in Figure 7).

**Figure 6.**
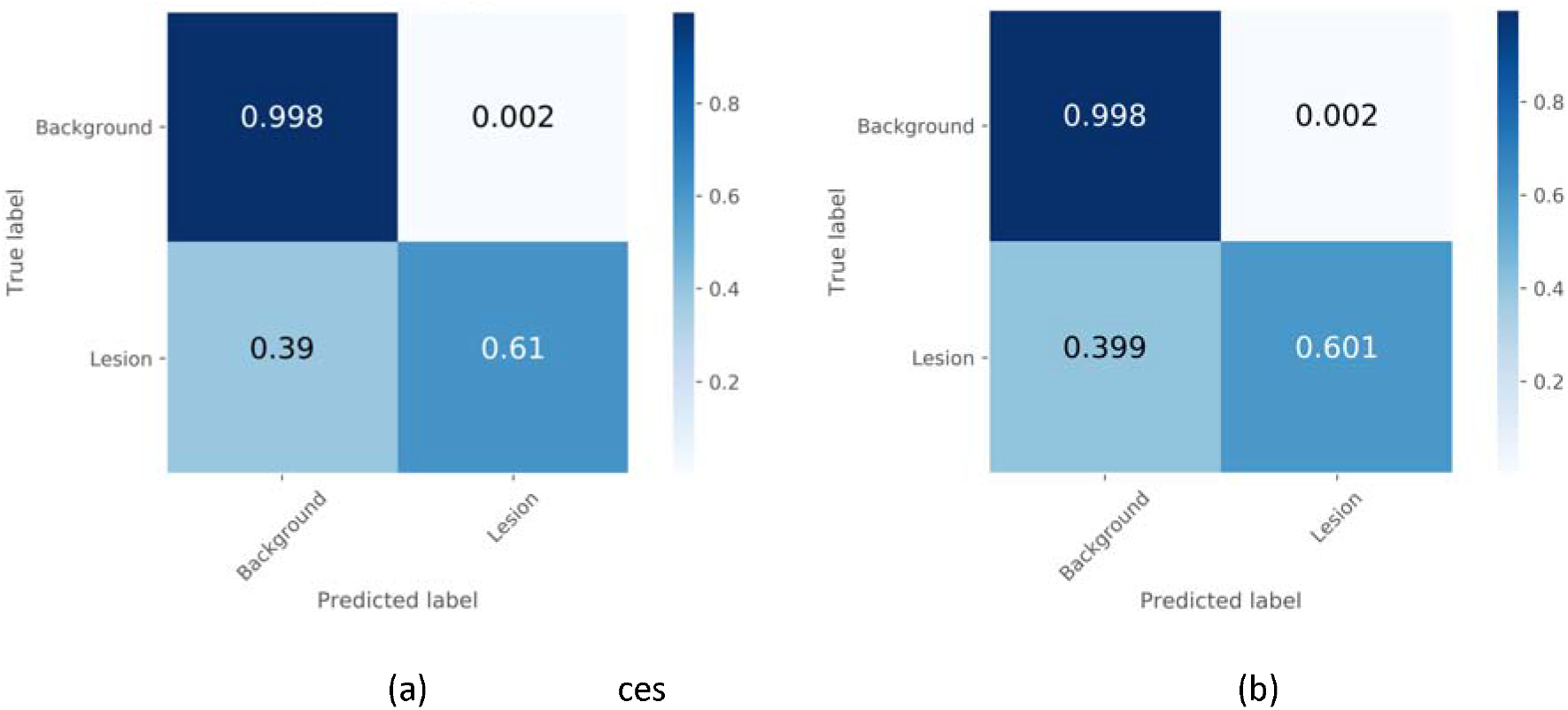
Confusion matrix in the validation dataset (a) and in the test dataset (b).

**Figure 7.**
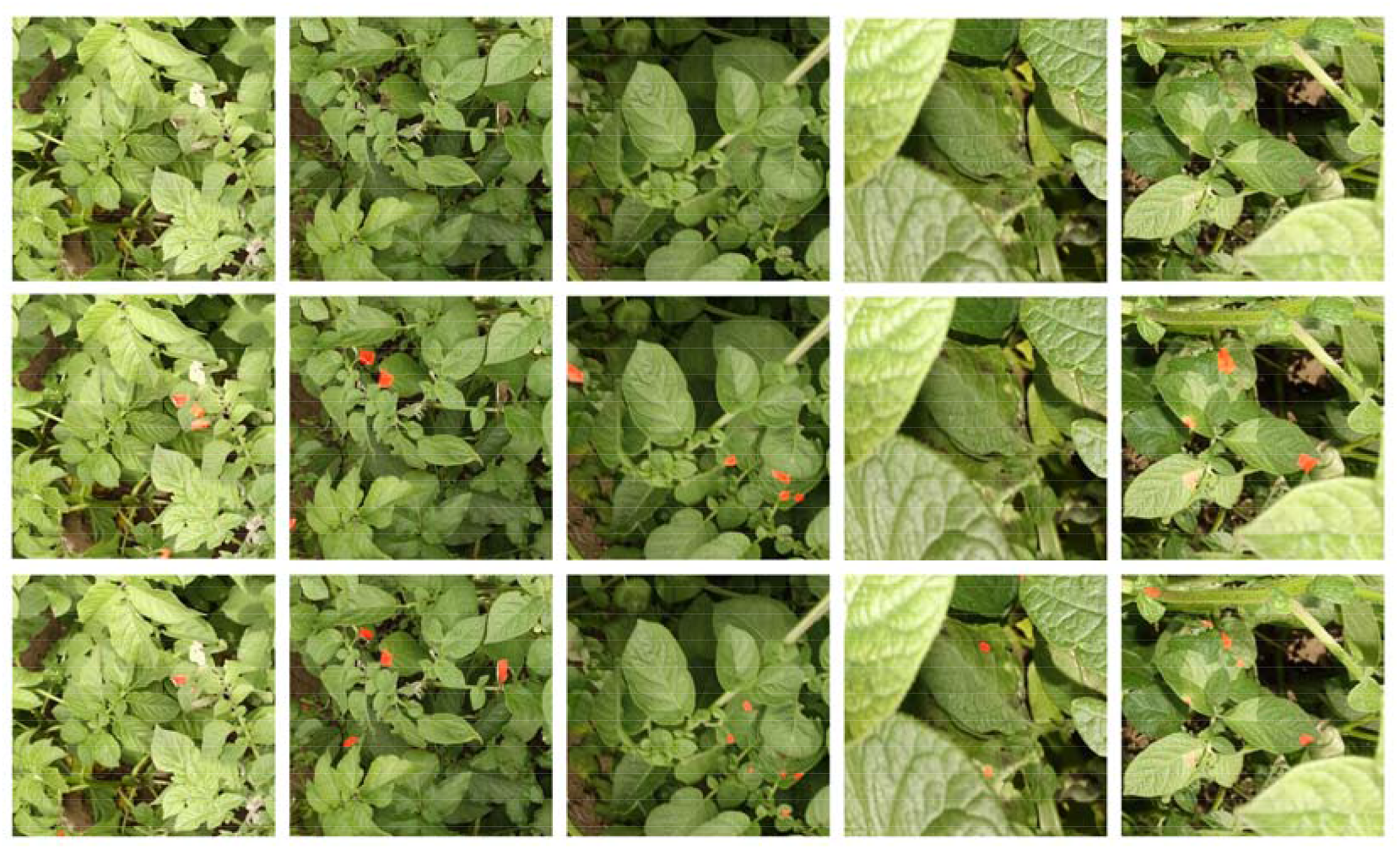
Examples of sub-image predictions (512×512) in the test dataset (row #1: raw sub-images, row #2: ground truth, row #3: predicted images)

The prediction masks of sub-images were reconstructed back to the predicted images of the original test images (5472×3648). Two examples of the predicted images are shown in Figure 8. There are some examples of failed cases in some predictions. For example, no disease lesions were visually observed on potato leaves (Figure 9). But 3 lesion areas were predicted by the model. The three false positives are all from soil patches which largely have similar shape features and areas as typical lesions. Moreover, these soil patches are surrounded by leaves, resulting in more confusion for inference. Also, some other wrongly predicted cases are located in the image border (#1 column in Figure 7). An entire lesion can be cut with two pieces when cropping a whole image into multiple sub-images. In this case, the partial lesion significantly changes the morphological features and loses the important neighbor pixel information for models to predict. This might lead to these failure cases.

**Figure 8.**
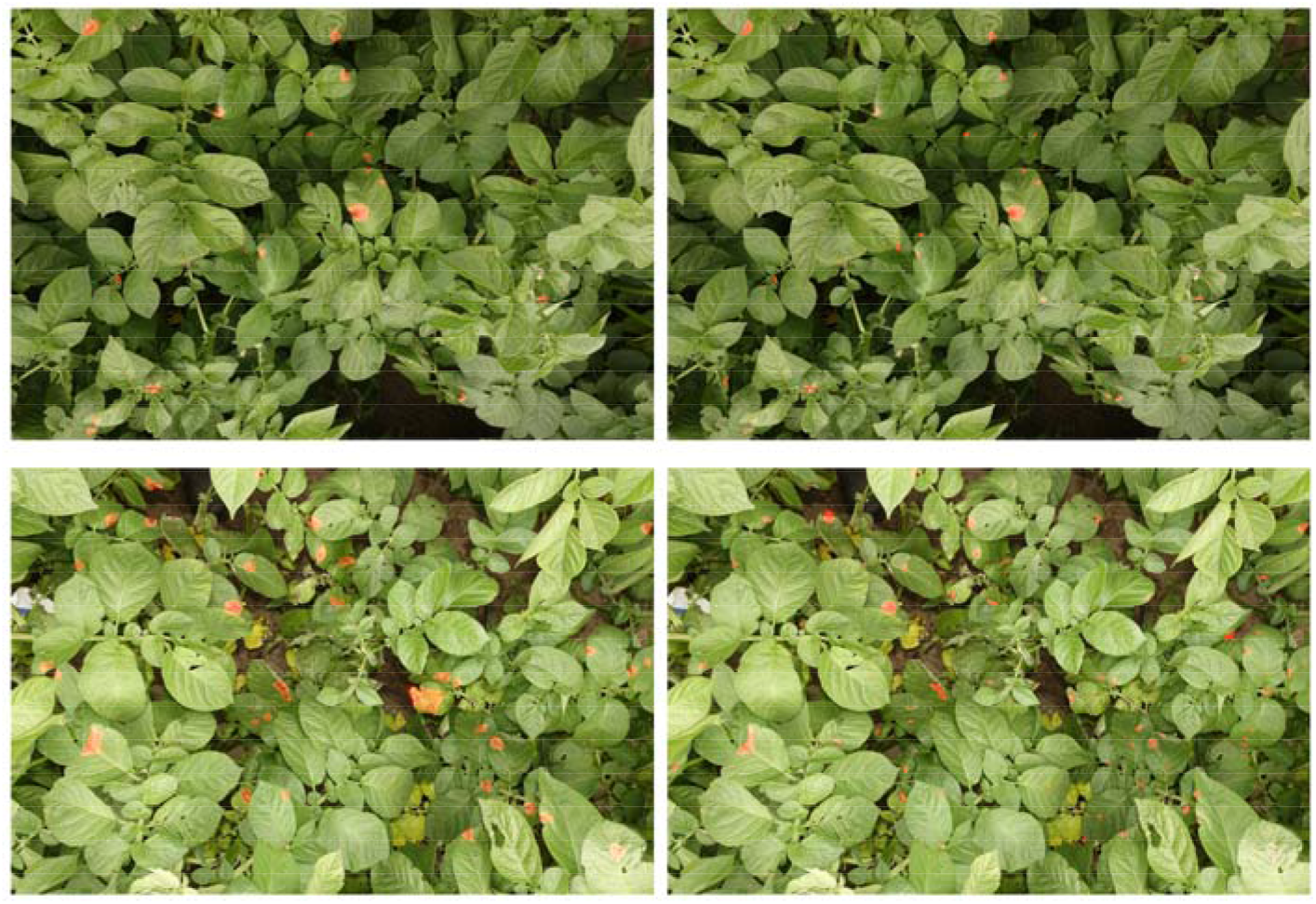
Examples of the predicted raw images (5472×3648) in the test dataset and ground truth images (Left column: ground truth images, Right column: predicted images)

**Figure 9.**
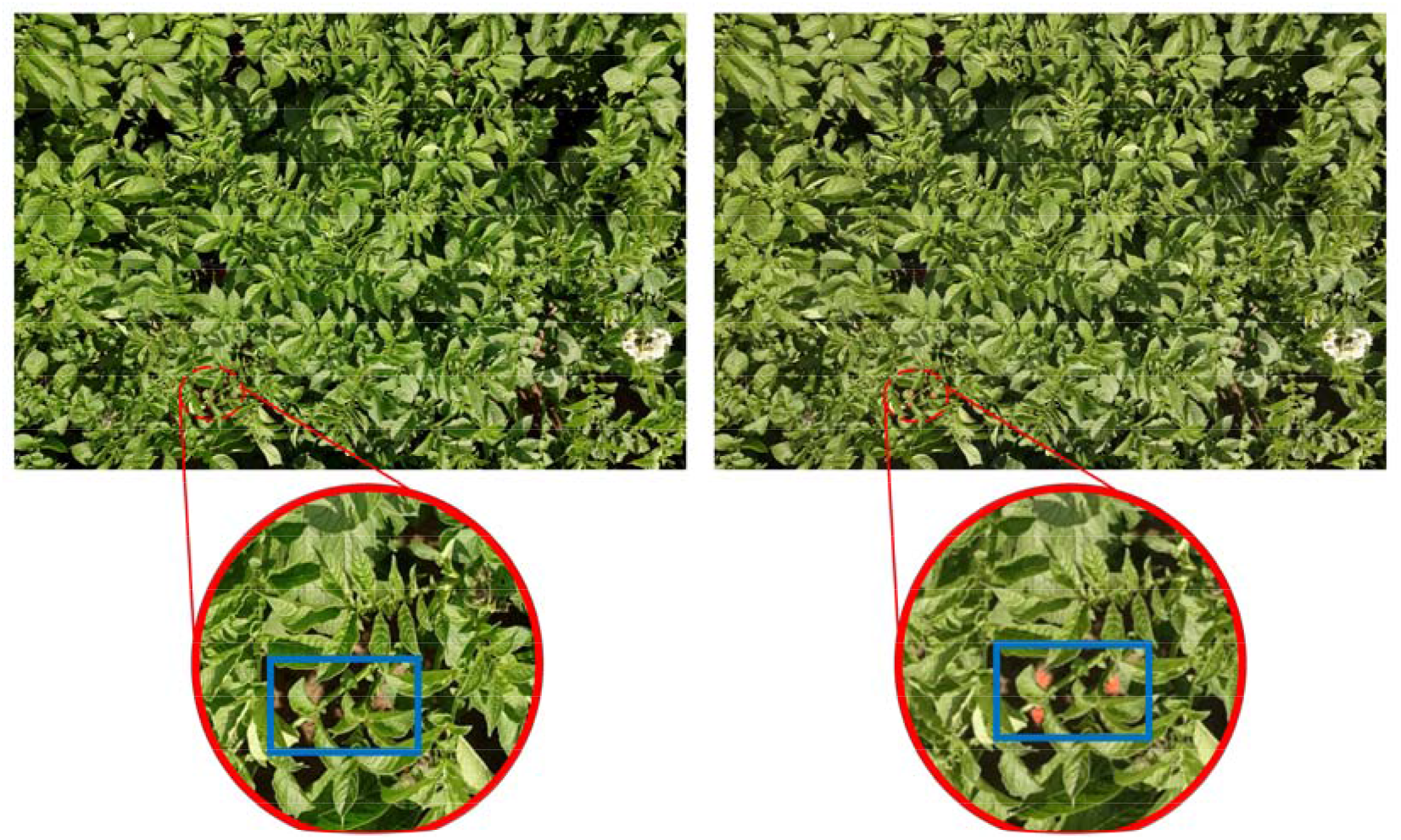
False positives in a test image (5472×3648).

### 3.4 Model validation in negative examples

We also tested the generalization ability and validity of the model with some difficult images (negative examples) without *P. infestans* lesions but with tissue damages caused by biotic or abiotic stresses. These images were collected from different potato fields in the summer of 2020. The model did not predict any lesions in these images in Figure 10. These damages were from various sources such as fertilization, herbicide and pathogens other than *P. infestans.* Figure 11 shows that the model failed to recognize the *P. infestans* lesions. Specifically, some lesions from *Alternaria solani* (Early Blight) were recognized as being *P. infestans* lesions. However, the model did not recognize damages in stems caused by leaf mold as *P. infestans* lesions, though they have similar color features as *P. infestans* lesions.

**Figure 10.**
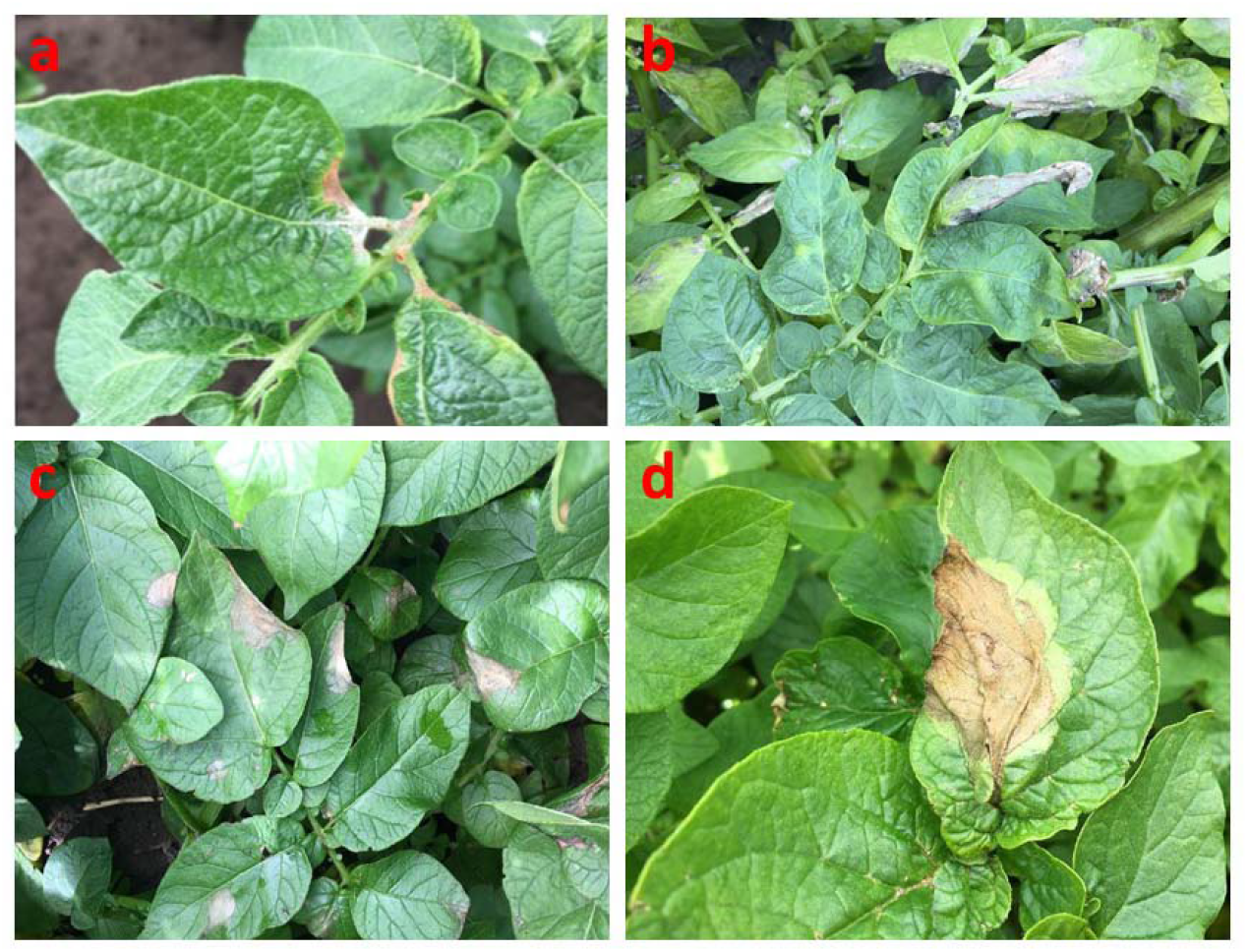
Correct prediction samples with damages of burning by lime nitrate (a); with deformity and necrosis caused by herbicide damage (b); with damage caused by eutrophication with lime nitrate (c); with infestation of possible grey mold (d).

**Figure 11.**
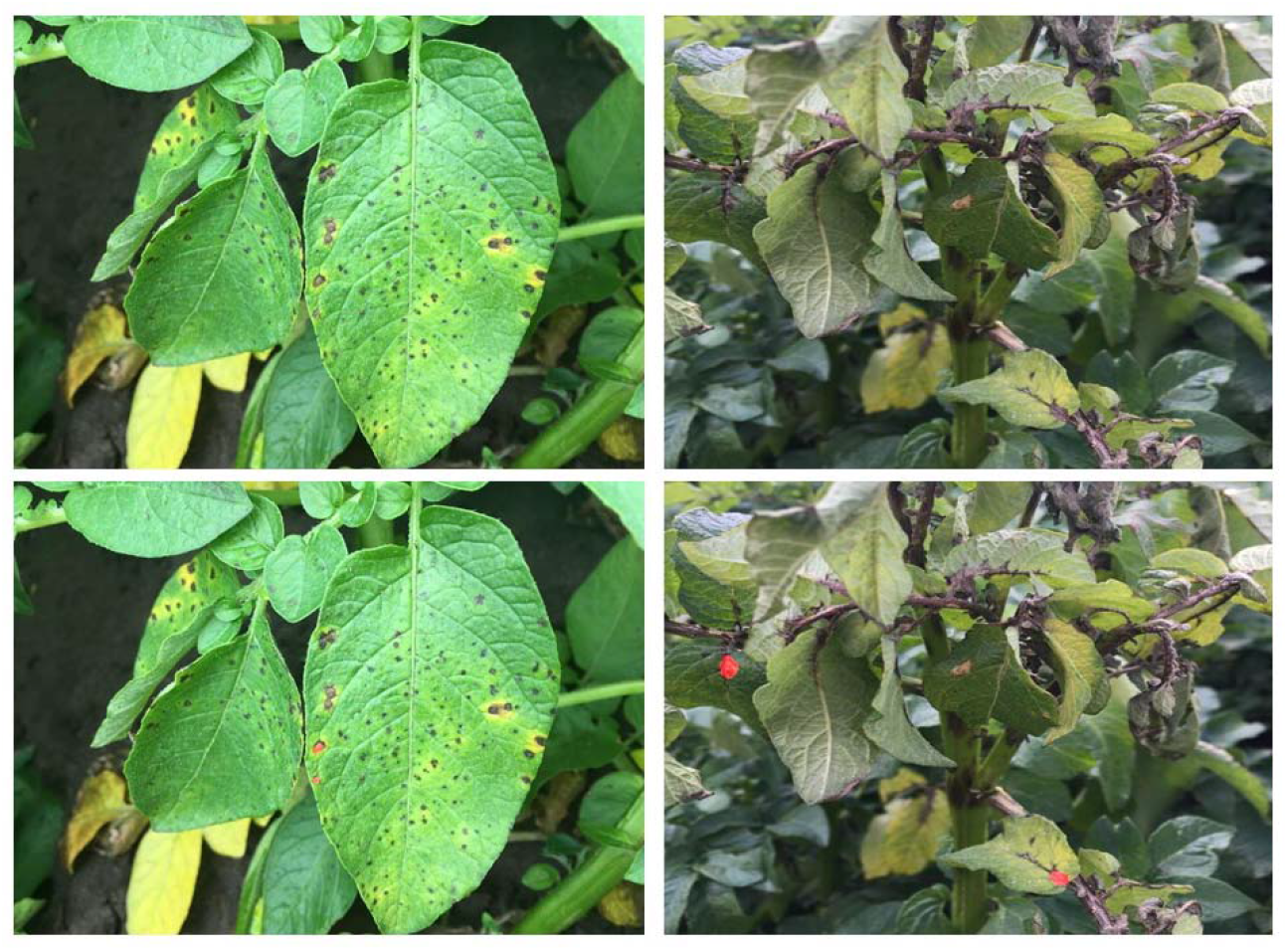
Failure cases of *P. infestans* lesion recognition where some cases of *Alternaria solani* infections are recognized (Left); and in an image with sever damages caused by leaf mold infestation (Right).

### 3.5 Correlation between visual scores and the number of lesions at the canopy level

As the visual scores were obtained based on the rating of the whole plant, we only selected the unseen images that covered the full crop canopy to avoid bias due to partial view. There were 43 images selected in total. The histogram of visual scores is shown in Figure 12. The average value of visual scores is 3% ranging from 0 to 40%. In order to minimize the failure cases raised from cropping, each image was predicted with multiple scales. Specifically, each image was cropped at 3×2, 4×3, 5×4, 6×5, 7×6, 8×7, 9×8 scales in horizontal x vertical directions. The number of lesions in each image was obtained based on the majority voting algorithm described in the section of post-processing. The histogram of the final detected lesions is displayed in Figure 13. There were 1063 lesions detected in these test images. The mean value of the detected lesion areas was 892 pixels. The maximum and minimum values are 7520 and 52 pixels, respectively. More than 40% of lesion areas are below 500 pixels, and very few lesion areas exceed 6000 pixels. This is consistent with the visual scores where around 80% of the scores are below 5%.

**Figure 11.**
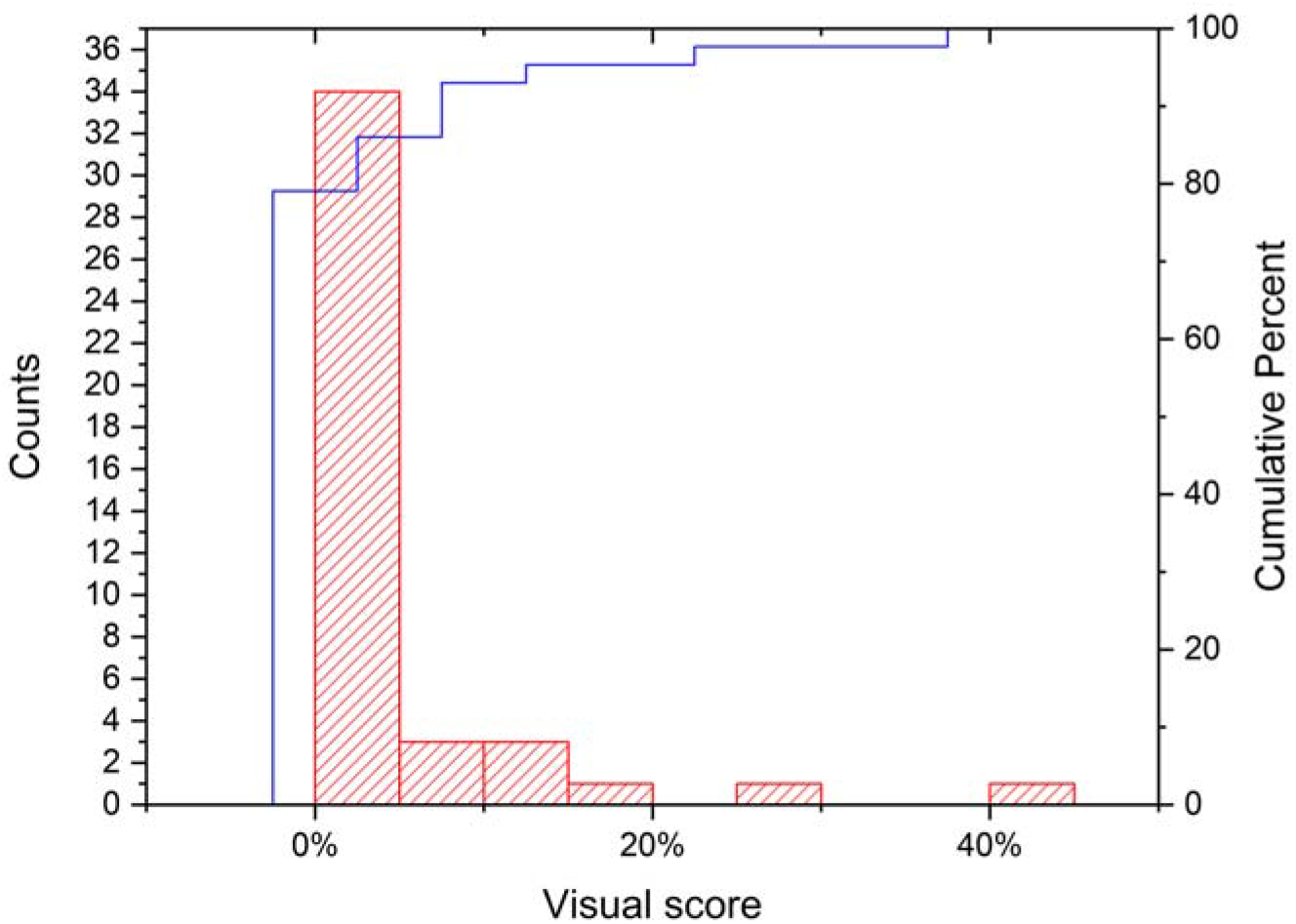
Histogram of visual scores; the red bar represents the number of visual scores in that x axis range (y axis in the left), and the blue line represents cumulative percentage of visual scores (y axis in the right).

**Figure 13.**
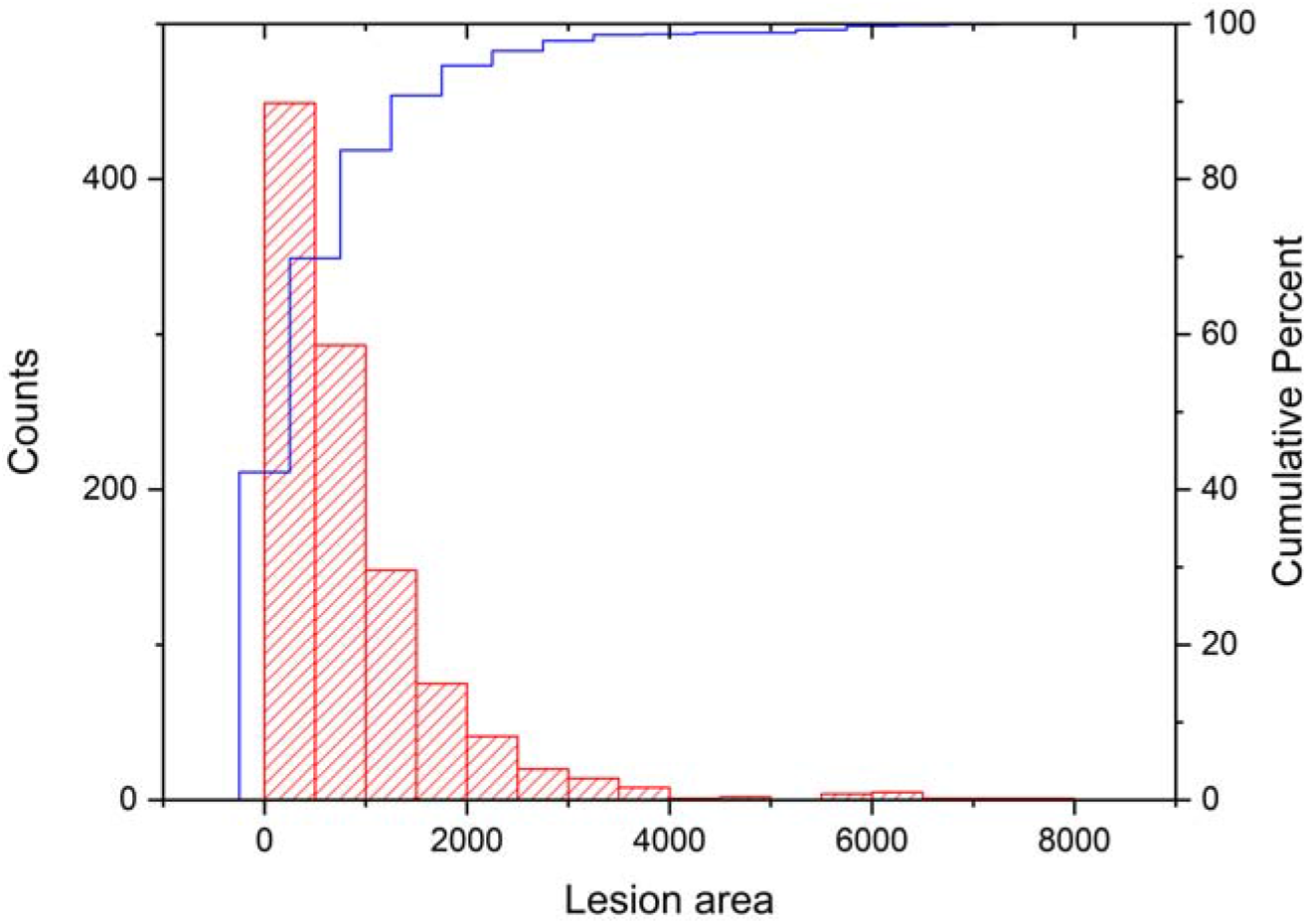
Histogram of lesion areas; the red bar represents the number of lesion areas in that x axis range (y axis in the left), and the blue line represents the cumulative percentage of lesion areas (y axis in the right).

A linear model was fitted, to quantify the relationship between visual scores obtained from an experienced plant breeder from the Danespo company and the number of lesions that appeared at canopy level. Figures 14 and 15 show the fitted linear relationships between visual scores and the number of lesions predicted from 3×2 and 5×4 scales, respectively. Figure 16 illustrates the fitted linear relationship between visual scores and the number of lesions obtained from majority voting of all scales. Compared to prediction masks with only one scale, the fused masks based on majority voting achieved a better linear relationship by increasing the R^2^ value from around 0.4 to 0.655.

**Figure 14.**
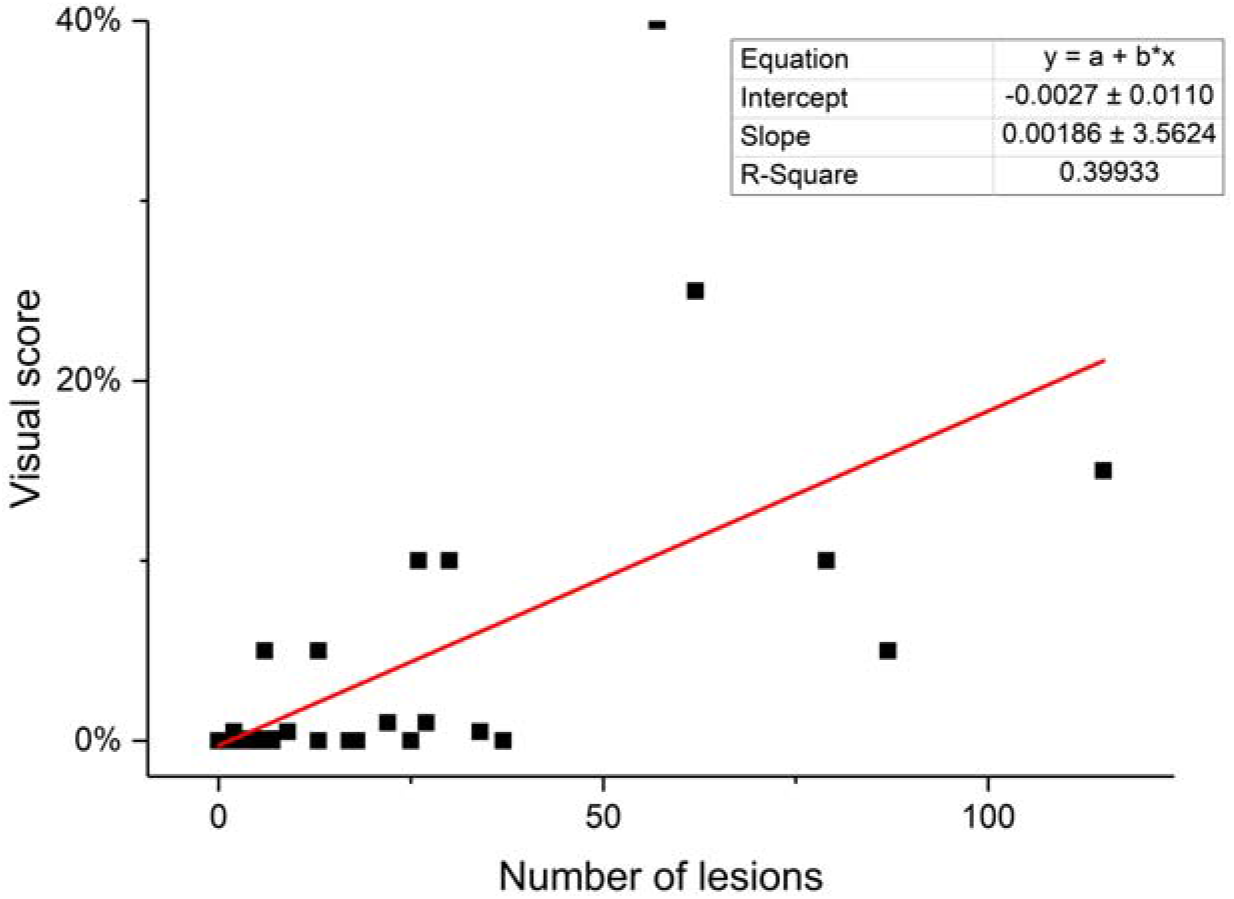
Fitted linear relationship between visual scores and number of lesions assigned from images from 3×2 scale.

**Figure 15.**
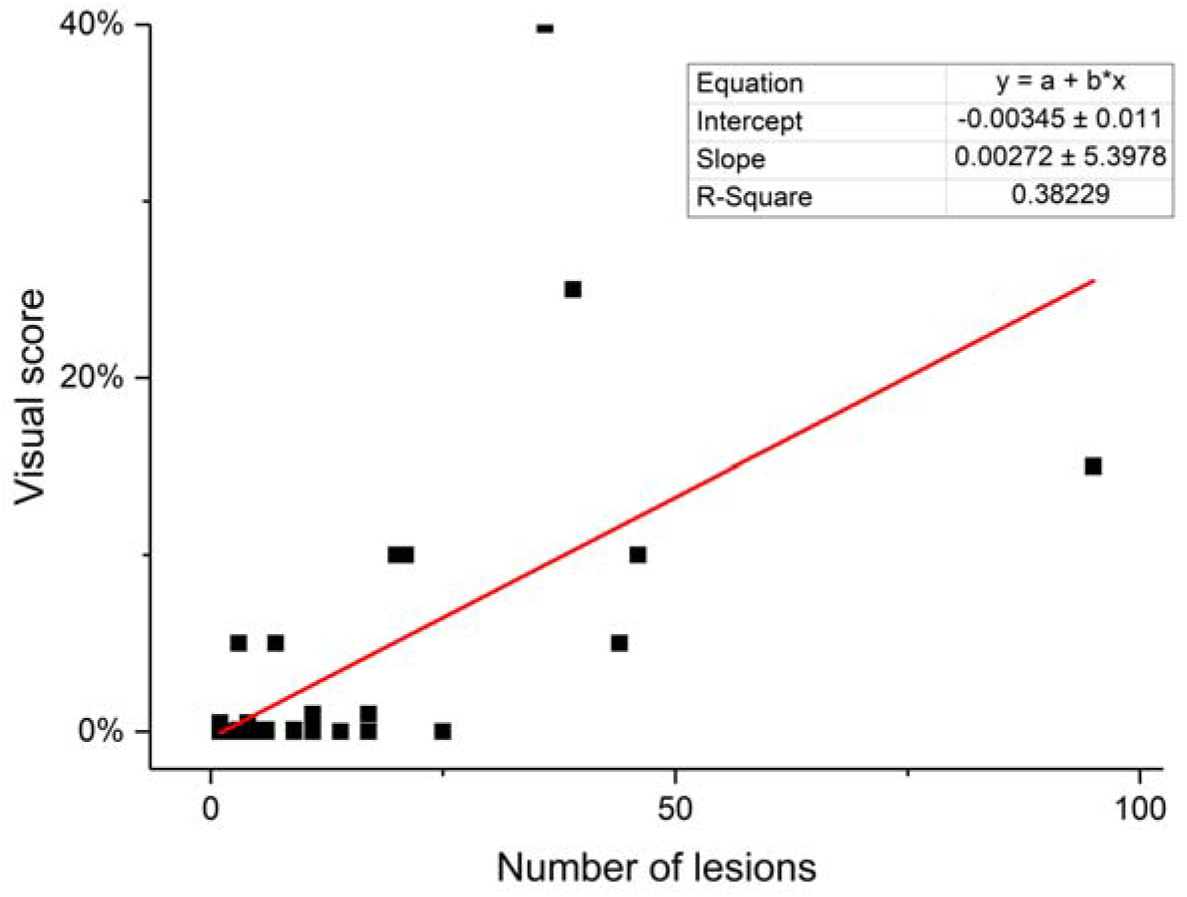
Fitted linear relationship between visual scores and number of lesions assigned from images from 6×5 scale.

**Figure 16.**
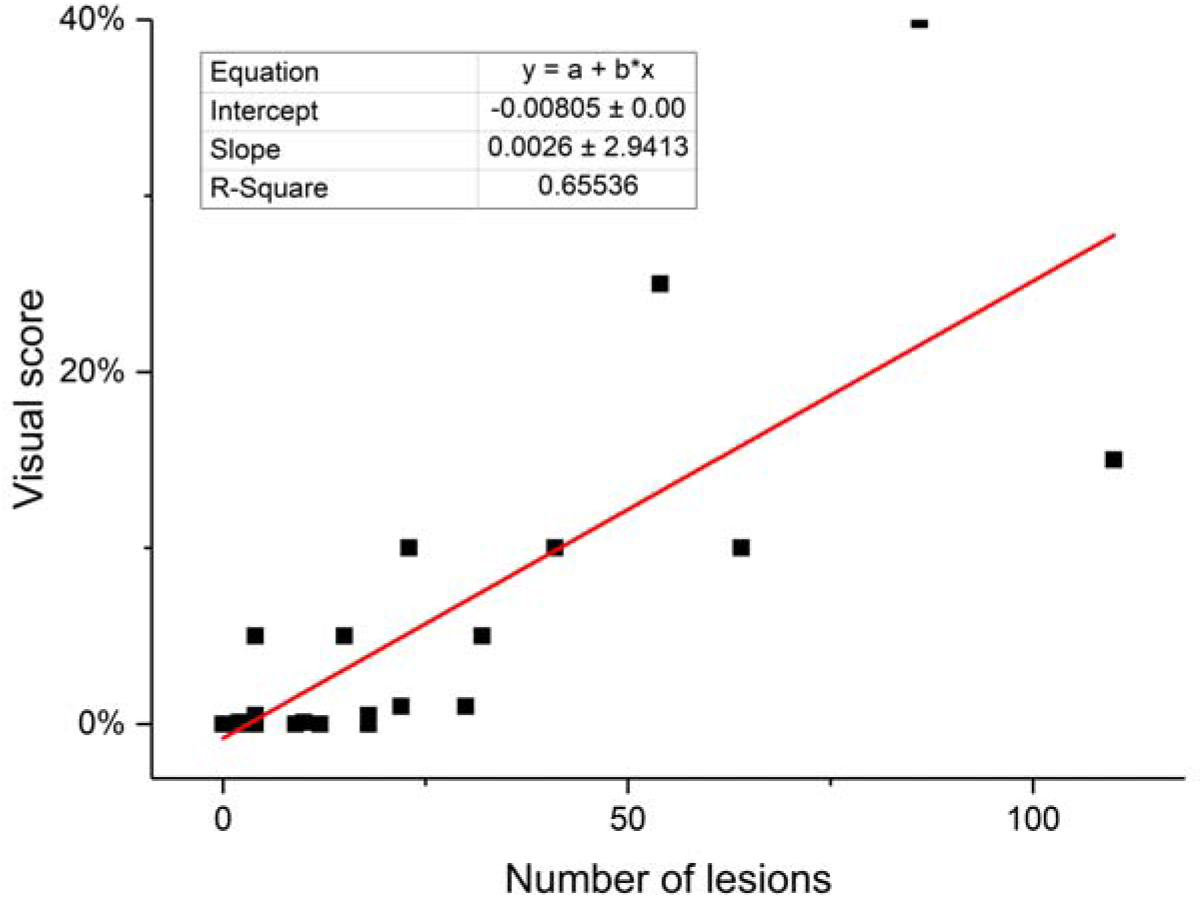
Fitted linear relationship between visual scores and number of lesions from majority voting from all scales.

## 4. Discussion

Deep learning has demonstrated its superior performance on disease detection in field settings compared to convolutional machine learning methods [17][20][18]. Imbalance classes are a common problem in deep learning and is widely represented in the agricultural field, and thus hampers applications in for example distinction between weed and crop [13] [30], pest detection [14] and disease segmentation [18]. In these cases, pixels of one class, generally soil or crop, are dominant in images. Training a network in an appropriate way is critical for delivering a good segmentation result for each class. Table 3 shows that assigning imbalance weights in the loss function can effectively mitigate the issue with imbalance classes, which is consistent with the conclusion by Yasrab et al (2019) [31]. The mIOU value is only 0.551 with same weights across classes, while it can be improved to nearly 0.7 with imbalance weight assignment. The IOU value (>0.99) of the background class (soil and crop plant) is far higher than the IOU value of disease lesion class. Based on the confusion matrix, it is concluded that majority of lesion pixels (>60%) were correctly classified. The relatively low IOU of lesion class (<0.5) indicates that a few of false positives (Figure 9) from soil patches, which have similar shapes to lesions were predicted by the model. Stewart et al [18] also found this kind of false positives for Northern Leaf Blight (NLB) lesion segmentation in maize fields. Besides, senesced leaves could also contribute to the false positives. Setting a threshold to filter out some false positives, based on lesion size, might improve the IOU value of lesion class. This threshold value can be estimated based on the prior knowledge of the maximum lesion area in certain developmental stage of PLB disease or plant development, related to the start of the outbreak or even disease prediction based on weather and cultivar. It should, however, be noted that a lesion size threshold is of course dependent on that images are collected at a fixed height. We drew a group of connected key points to define the lesion area when manually labelling images with the tool LabelMe. This way of annotating speeds up the pixel-wise labelling process for segmentation. However, it is difficult to always draw a very accurate boundary line with those key points especially since a majority of PLB lesions in our images are tiny and have an irregular shape. As a consequence, the labelled lesion areas at times inevitably include pixels belonging to the background class (soil and leaf), leading to the relatively low IOU of lesion class. To overcome this problem, Wiesner-Hanks et al [19] discussed using crowdsourced data for NLB lesion detection at millimeter-level based on aerial visual images and concluded that increasing the number of workers per image could improve the quality of annotation polygons.

Image preprocessing is essential before feeding images to train a neural network. It is encouraged to randomly apply blur, contrast and brightness as data augmentation for the benefit of model robustness. The training dataset with 1600 labelled images from 31 potato genotypes is still limited and unlikely to include all lesion variations. We tested the generalization ability and validity of model in recognition of *P. infestans* lesions with some images from other fields and conditions (Figure 10). The failure cases featured highly similar colour and morphological characteristics as *P. infestans* lesions still are represented. Including such failed images in the training dataset again can improve the performance [32]. The creation of further lesion variations in the training datasets by synthetic lesion images could be a good way to improve model performances. To this end, Sun et al [33] developed a conventional image processing algorithm to optimize synthetic lesion images obtained from a generative adversarial network (GAN). Cap et al [34] proposed a LeafGAN algorithm for lesion image generations, which improved the diagnostic performance by 7.4%.

The use of a majority voting to generate accurate lesion masks from multiple prediction scales, inspired from random forest machine learning algorithm described in [35], proved its effectiveness to establish a linear relationship between visual scores and number of lesions at canopy level (Figure 16). Very few studies have tried to automatize the visual scoring in field environments for plant breeding. In reality, visual scores are evaluated based on the number of lesions and their areas on single leaflets at early infection stages, which brings difficulties with analysis based on with 2D images at the canopy level. The lesion recognition should be further explored by three-dimensional (3D) imaging to obtain full plant structures and by employing the state-of-the-art network architecture in semantic segmentation to reduce the inference time.

In precision farming, it is necessary to detect PLB disease as early as possible to bring in appropriate measurements to avoid yield loss. As only RGB images were used in this analysis, pre-symptomatic detection of PLB is inevitable missed. For pre-symptomatic crop disease detection, hyperspectral measurements from spectroradiometers or spectral imaging sensors are generally employed [36]. For example, Anderegg et al [37] used an ASD FieldSpec spectroradiometer (350-2500nm) to measure wheat plants at a canopy level for *Septoria Tritici* Blotch (STB) disease detection and quantification. Gold et al [38] measured contact leaf reflectance with a field spectrometer for pre-symptomatic PLB detection. For many applications [39] [40] [41], spectral imaging sensors are more popular than non-imaging hyperspectral sensors due to the additional capability of providing spatial information on shape, texture and color. Partial least square discriminant analysis (PLSDA) is generally used to process full spectral data [42], Our study also has the potential to monitor the development of PLB disease after lesion appearance, which could be useful to screen for high PLB resistance potato genotypes from a diverse germplasm in precision breeding. In terms of PLB management in potato production, it is useful to acknowledge how early PLB should be detected after the appearance of lesions. To address this question, Wiik et al [43] carried out trials over two years in 2018 and 2019 to test the need of first intervention after the first visual symptom were detected. The preliminary results showed that it is acceptable for farmers to apply a first spray with curative systematic fungicides after the discovery of first symptom, corresponding to a very low 0.01% infection provided that it is sprayed more or less immeadiatly as it was shown that a first spray delayed by 5 days later first symptom appears was too late to stop the disease. 0.01% infection corresponds to to only 300 spots/ha if the disease is evenly spread over the field. Thus it is clear that protection against PLB for precision agriculture requires very early detection of the symptoms with RGB.

## 5. Conclusion and future work

In this study, we demonstrated the feasibility of using a deep learning algorithm based on an encoder-decoder architecture for potato late blight disease lesion semantic segmentation based on field images. The results show that the intersection over union (IoU) values of background (soil and leaf) and lesion classes in the test dataset are 0.996 and 0.386, respectively. Assigning different weights for imbalance class could improve the performance of the model. This work also presents the possibility of accurate lesion counting at the plant canopy level with the use of image alignment. A linear relationship between visual scoring and the number of lesions was established. We can also conclude that the fused masks obtained from majority voting of the masks predicted with multiple scales achieved higher R^2^ value (0.655) compared to prediction with a single scale. The proposed methodology has the potential to monitor the lesion development under field conditions and evaluate the resistance of genotypes against potato late blight enabling more precise and automated potato breeding.

This study will be followed by further field tests and the model will continue to be tested in terms of robustness and accuracy by adding new field image datasets. The updated model will also be used to test on images collected from different time points to predict area under disease progression curve (AUDPC). In addition, we will continuously update the models with the new labeled datasets and synthetic images to improve the generalization ability. Multiple imaging sensors like multispectral and hyperspectral cameras hold promise to also detect and maybe even quantify pre-symptomatic disease. Also, the sensor combinations, e.g. a spectroradiometer (early stage) and high-resolution RGB camera (late stage), can be considered for monitoring PLB progression. Multimodal data fusion and machine learning are suggested to be fully exploited for this application. Furthermore, it would be interesting to explore new vehicle and sensor techniques to build three-dimensional imaging to be able to detect disease lesions below the canopy.

## Acknowledgement

We acknowledge the Flemish Supercomputer Center (VSC) for providing the GPU computational resources and services for this work. We thank Mathieu Gremillet for field assistance, Hanne Grethe Kirk at Danespo for visual scoring of disease, and Linnea Almqvist from SLU for providing image examples in Figures 10 and 11. This research was founded by Nordic Council of Ministers (PPP #6P2), NordForsk (#84597) and Vinnova (#2016-04386).

## Notes

### Competing Interest Statement

The authors have declared no competing interest.

